# Interferon-gamma is Quintessential for NOS2 and COX2 Expression in ER^-^ Breast Tumors that Lead to Poor Outcome

**DOI:** 10.1101/2023.04.06.535916

**Authors:** Robert YS. Cheng, Lisa A. Ridnour, Adelaide L. Wink, Ana L. Gonzalez, Elise L. Femino, Helene Rittscher, Veena Somasundarum, William F. Heinz, Leandro Coutinho, M. Cristina Rangel, Elijah F. Edmondson, Donna Butcher, Robert J. Kinders, Xiaoxian Li, Stephen T.C. Wong, Daniel W. McVicar, Steven K. Anderson, Milind Pore, Stephen M. Hewitt, Timothy R. Billiar, Sharon Glynn, Jenny C. Chang, Stephen J. Lockett, Stefan Ambs, David A. Wink

## Abstract

A strong correlation between NOS2 and COX2 tumor expression and poor clinical outcomes in ER-breast cancer has been established. However, mechanisms of tumor induction of these enzymes are unclear. Analysis of The Cancer Genome Atlas (TCGA) revealed correlations between NOS2 and COX2 expression and Th1 cytokines. Herein, single cell RNAseq analysis of TNBC cells shows potent NOS2 and COX2 induction by IFNγ combined with IL1β or TNFα. Given that IFNγ is secreted by cytolytic lymphocytes, which improve clinical outcomes, this role of IFNγpresents a dichotomy. To explore this conundrum, tumor NOS2, COX2, and CD8^+^ T cells were spatially analyzed in aggressive ER-, TNBC, and HER2+ breast tumors. High expression and clustering of NOS2-expressing tumor cells occurred at the tumor/stroma interface in the presence of stroma-restricted CD8^+^ T cells. High expression and clustering of COX2-expressing tumor cells extended into immune desert regions in the tumor core where CD8^+^ T cell penetration was limited or absent. Moreover, high NOS2-expressing tumor cells were proximal to areas with increased satellitosis suggestive of cell clusters with a higher metastatic potential. Further *in vitro* experiments revealed that IFNγ+IL1β/TNFα increased elongation and migration of treated tumor cells. This spatial analysis of the tumor microenvironment provides important insight of distinct neighborhoods where stroma-restricted CD8^+^ T cells exist proximal to NOS2-expressing tumor niches that could have increased metastatic potential.

## Introduction

Estrogen receptor alpha-negative (ER-) and triple negative breast cancer (TNBC) account for a smaller proportion of breast cancer types but are among the most aggressive malignancies with limited treatment strategies when compared to less aggressive ER+ tumors (1). During the past decade, a significant proportion of cancers have demonstrated elevated NOS2 expression (2), where cancers ranging from melanoma to glioma overexpress NOS2 (3–6). In breast cancer, increased NOS2 has been reported in >70% of patients (7). Interestingly, elevated tumor NOS2 expression correlated with P53 mutation and was predictive of poor survival in ER- but not ER+ breast cancer patients (7, 8). While elevated tumor COX2 expression was also predictive of poor outcome as defined by a Hazard Ratio (HR) of 2.45 in ER-patients from the same cohort (9), elevated NOS2/COX2 co-expression was strongly predictive of poor outcome (HR 21) (10). These results suggest that elevated tumor NOS2/COX2 coexpression drive the progression of aggressive breast cancer phenotypes (10, 11), however mechanisms of NOS2/COX2 induction within tumors remain unclear.

Examination of The Cancer Genome Atlas (TCGA) has revealed correlations between tumor NOS2/COX2 expression and interferon-gamma (IFNγ), interleukin-17 (IL17), IL1, and toll-like receptor-4 (TLR4), which are frequently associated with anticancer effects (12). Interestingly, these associations are contradictory as elevated tumor NOS2/COX2 expression predict poor clinical outcomes (7, 9, 10). Recently, elevated NOS2 and COX2 expression was discovered in distinct immune and tumor cells upon treatment with IFNγand cytokines or TLR4 agonists, consistent with feed forward NOS2/COX2 signaling as previously reported (10, 13). These results suggest an orthogonal relationship between tumor NOS2/COX2 expression, which could promote distinct tumor microenvironments that contribute to poor clinical outcomes (13).

To explore these possibilities, herein we show that Th1 cytokines effectively stimulate NOS2 and COX2 expression in tumor cells in vitro. Single-cell RNAseq (scRNAseq) analysis revealed that IFNγ combined with interleukin 1β (IL1β), or tumor necrosis factor alpha (TNFα) induced higher NOS2/COX2 expression than the cytokines as single agents. In addition, cytokines that were upregulated in high NOS2-expressing ER-breast tumors (7, 10), including IL6 and IL8, were induced by these treatments and correlated with NOS2/COX2 expression. Also, this study reveals a unique synergy between IL1α/β that enhances the expression of NOS2 and COX2. Multiplex spatial imaging revealed clusters of high NOS2 expressing cells proximal to areas of stroma restricted CD8^+^ T cells that are known to produce IFNγ and suggests a small inflammatory niche at the tumor/stroma interface in these tumors. While COX2 was present in these regions, it was more highly expressed further into the tumor, in immune desert areas with low CD8^+^ T cell penetration. Comparison of distinct sites within the same tumor, or geographic regions between tumors, suggests a spatial and temporal progression of inflammatory sites in areas of restricted lymphoid cells, which progresses to an immune desert in areas of high COX2 expression. These novel observations provide distinct spatial fingerprints of aggressive tumor phenotypes that correlate with decreased disease-specific breast cancer survival (7, 10).

## Results

### TCGA and Bioinformatics for Tumor NOS2/COX2 Expression

Our previous work comparing high and low NOS2 and COX2 (also known as PTGS2) tumor expression revealed associations with a variety of inflammatory markers that are typically associated with antitumor activity (14), which prompted us to explore cytokine regulatory effects on NOS2/COX2 expression in tumor cells and tissues. A correlation analysis was performed using the Xena browser and TCGA database to gain a deeper understanding of the conditions associated with NOS2 and COX2 expressions in ER- breast cancer. The correlation analysis of TCGA-BRCA (breast cancer) revealed several inflammatory pathways associated with increased tumor NOS2/COX2 expression (Figure 1A) that influenced clinical outcomes (Figure 1B), including the antitumor-associated IFNγ, TNFα, IL2, IL1, and IL17 pathways (Figure 1A). Moreover, tumor NOS2/COX2 expressions were positively correlated and Th1 cytokines had the strongest positive association with COX2, whereas IL1β was associated with NOS2 (Figure 1A). These findings suggest a dichotomy within the tumor microenvironment (TME), where cytokines that are generally associated with favorable outcomes also promote elevated tumor NOS2/COX2 expressions (Figure 1A), which are strong predictors of poor disease-specific survival in ER-breast cancer patients (Figure 1B) (7, 9, 10). To confirm the regulatory roles of these cytokine(s) in the induction of NOS2 and/or COX2 expressions, MDA-MB231 (MB231) breast cancer cells were stimulated with IFNγ in the presence and absence of TNFα, IL1β, IL17, and the TLR4 agonist lipopolysaccharide (LPS). Single-cell RNAseq data revealed that 48 hr exposure to IFNγ combined with TNFα or IL1β induced the highest expressions of NOS2 and COX2 (Figures 1C). Consistent with previous reports in murine tumors (12), high NOS2 expression requires IFNγ and TNFα/IL1, which exhibit a striking difference in cytokines produced during induction of murine macrophages and tumor cells (12). Moreover, scRNAseq of cytokine stimulated MB231 cells revealed that a 48hr stimulation with IFNγ in the presence of IL1β or TNFα induced the highest NOS2 and COX2 expressions, which clustered in the same regions of the t-SNE plot (Figure 1C and 1D). These results confirm high NOS2 and COX2 expression is induced by IFNγ combined with IL1β or TNFα as shown in the TCGA analysis. As predicted, scRNAseq analysis of IFNγ + IL1β/TNFα revealed an increase in the transcript levels of NOS2 and COX2 in treated MB231 cells. The number of NOS2 transcripts ranged from 1 to 3, whereas COX2 transcripts had a significantly wider range (Figure 1D). While less than 9% of the cells contained NOS2 transcripts, up to 40% of the cells contained COX2 transcripts (Figure 1D). NOS2 expression required IFNγ, whereas COX2 expression was weakly induced by IL1β or TNFα alone (Figure 1D). Nonetheless, in a manner similar to NOS2, the strongest COX2 expression occurred by IFNγ+IL1β/TNFα. In addition to cytokine stimulation, NOS2/COX2 feed forward signaling could also promote elevated NOS2/COX2 expression (Figure 1A) as previously reported (14).

**Figure 1.**
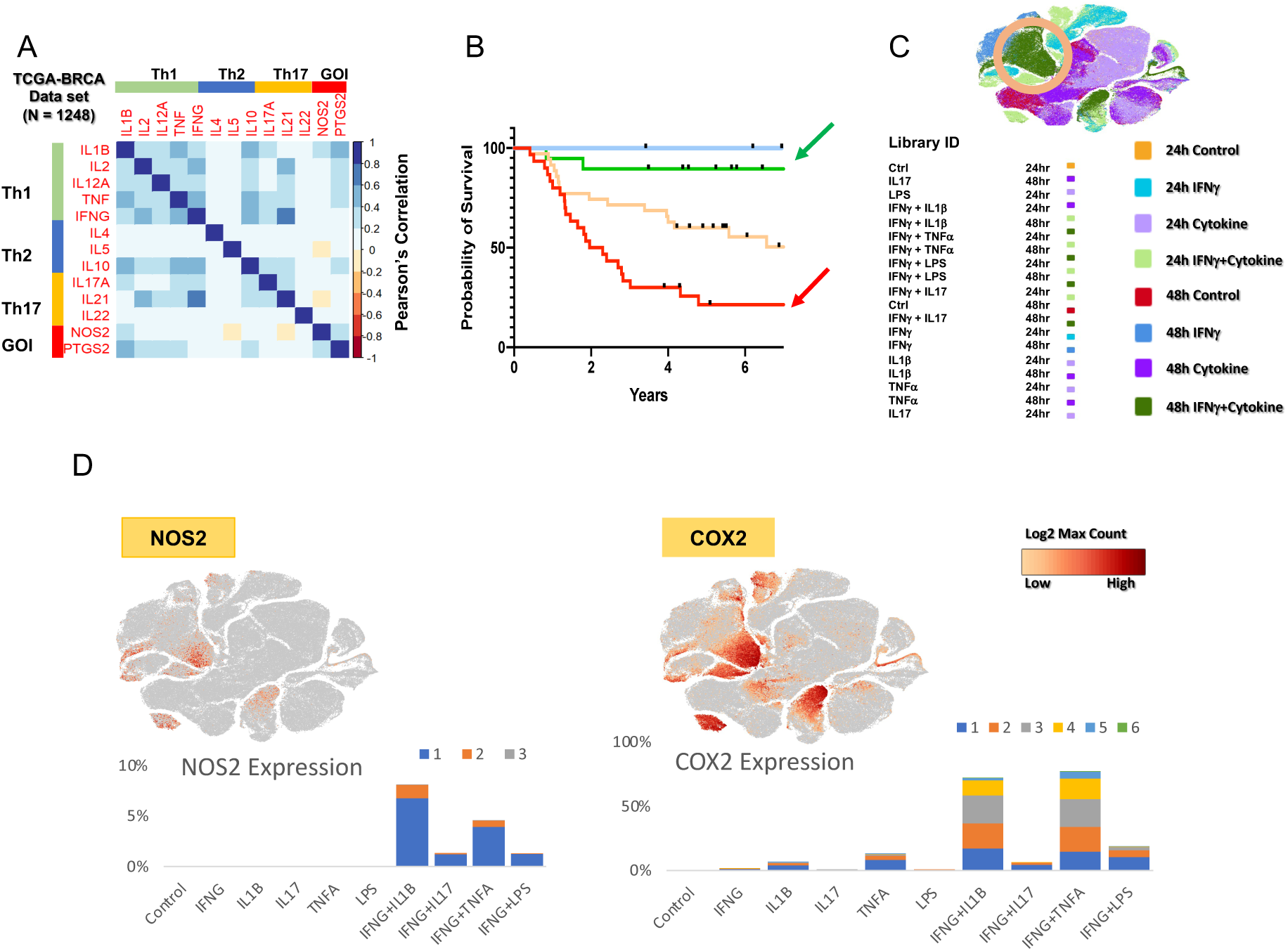
IFNγ and cytokines induce maximal expression of NOS2 and COX2 in ER- breast cancer cells. A) A heatmap display of Pearson’s correlation analysis of TCGA-BRCA (n = 1248) database through the UCSC Xena browser analyzing Th1, Th2, Th17 and two GOI (gene of interest) genes. The heatmap was generated in corrplot (0.92) in R (4.2.1). B) Survival analysis associated with NOS2_lo_/COX2_lo_ (green arrow) and NOS2_hi_/COX2_hi_ (red arrow) tumor protein expressions. C) t-SNE plot (Loupe Browser 6.3.0) of single cell analysis of MB231 cells treated with single or combination of cytokines IFNγ (100U/ml), IL1β(10 ng/ml), TNFα (10 ng/ml), IL17 (100ng/ml) and LPS (10 ng/ml) for 24 and 48 hr. Light and dark green color clusters represent 24- and 48-hr time points associated with IFNγ + IL1β/TNFα treatment, respectively. The orange circle indicates the highest overlapped cell clustering for the IFNγ+IL1β or TNFαafter 48 hr treatment. D) t-SNE plots of NOS2 and COX2 clustering cells. Stacked bar charts show the number of NOS2 and COX2 transcripts per cell in 48 hr treatment groups. Transcript per cell data (color code: blue, 1; orange, 2; grey, 3; yellow, 4; cyan, 5; green, 6) were extracted using the R packages (data.table 1.14.2, dplyr 1.0.10, and ggplot2 3.3.6).

Cancer stemness markers have been reported in cell populations with elevated CD44 and reduced CD24 expression levels that exhibited the same drug-resistant histopathological features of the derived tumor when injected in mice at very low concentrations (15). Herein, t-SNE plot analysis of CD44/CD24 expression revealed distinct clustering patterns (Figure 2A) defined by increased CD44 and reduced CD24 levels in high NOS2/COX2 expressing clusters after 48 hr treatment with IFNγ + IL1β or TNFα. This pattern is suggestive of increased cancer stemness (15) and is consistent with earlier reports of elevated CD44 levels in high NOS2 expressing ER- breast tumors (7, 16). The expression of tissue inhibitor of metalloproteinase-1 (TIMP1), a fibrosis marker (17), was similar to CD44, indicating that their expressions were likely controlled by the same upstream regulator (Figure 2A). To explore the expression relationships between Th1 cytokines, we analyzed the clustering patterns of IL6, IL8, IL1α, and IL1β (Figure 2B). The clustering patterns of these cytokines are highly similar and strongly overlap after 48 hr stimulation with IFNγ + IL1β/TNFα (Figure 2B). Moreover, examination of other cytokine-induced genes revealed a strong association with IL1α/β (Figure 2C) and are consistent with observations demonstrating that circulating IL1β predicts poor survival (18). Together, these findings support the TCGA-BRCA correlation analysis shown in Figure 1A.

**Figure 2.**
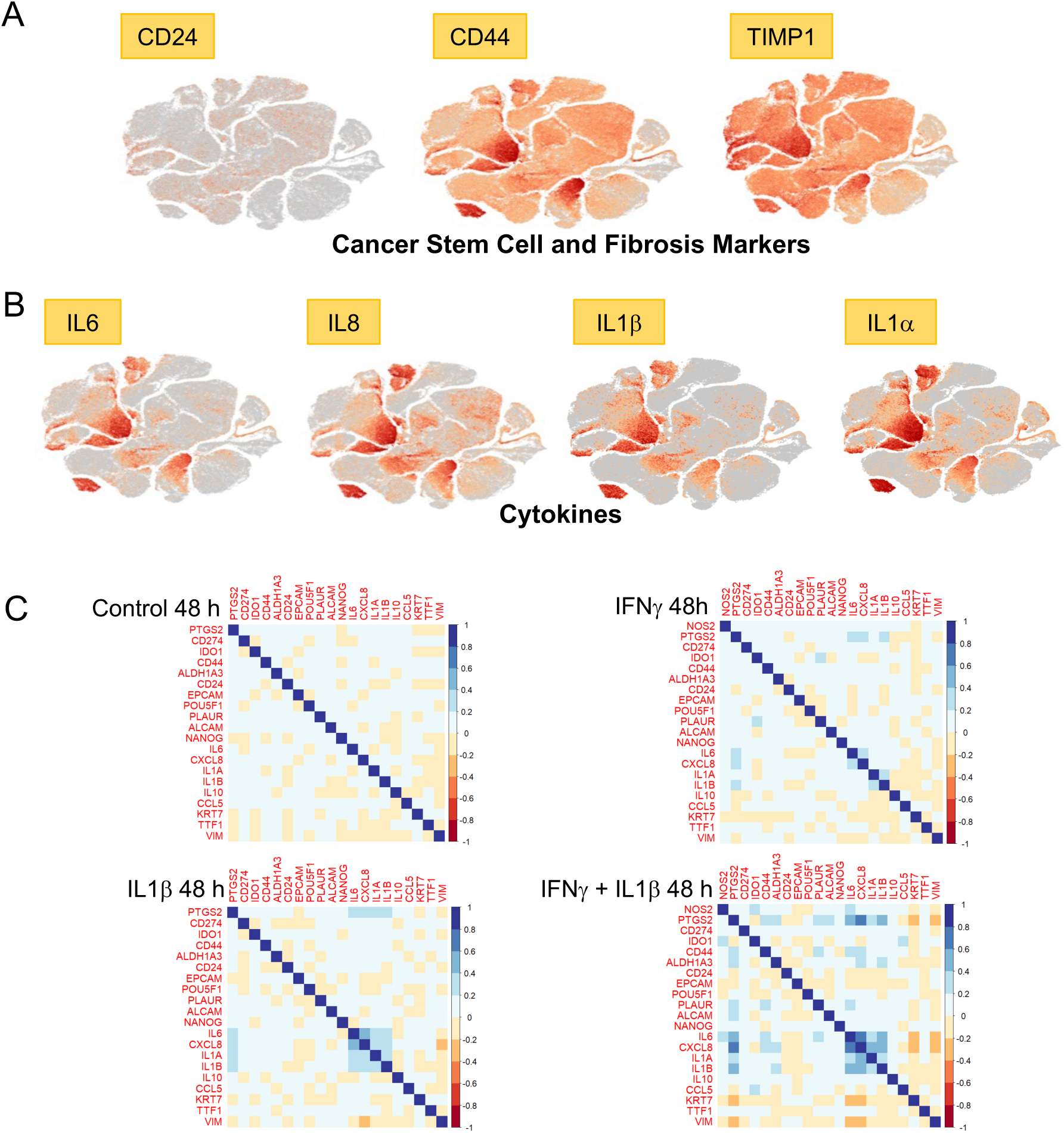
t-SNE plot and correlation analyses. A) t-SNE plots of cancer stem cell (CD24/CD44) and fibrosis (TIMP1) markers and B) Th1 cytokine (IL6, IL8, IL1β, IL1α) markers, which clustered in the same regions as NOS2/COX2 shown in panel A. C) Correlation analysis of selected cytokines, cancer stem cell markers, and metastatic-associated genes. Data were extracted from 48 hr Control, IFNγ, IL1βand IFNγ+IL1β treated groups in corrplot (0.92) in R (4.2.1). NOS2 was not expressed in Control and IL1β treated cells and is therefore not represented in the correlation analysis.

### Spatial Identification of NOS2 and COX2 Niches

The above *in vitro* findings demonstrate that IFNγ is a key regulator in the induction of NOS2/COX2 expression in ER- breast tumor cells, which raises the question of the origin of IFNγ secretion in tumor tissues. Cytotoxic lymphocytes release IFNγ and are associated with improved survival in TNBC and other cancer types (19–21). In contrast, elevated tumor NOS2/COX2 expression promote disease progression and are strongly predictive of poor disease-specific breast cancer survival (7, 9, 14). These findings implicate a dichotomy where anti-tumor lymphoid-producing IFNγ cells could induce pro-tumor NOS2/COX2 expressing cellular niches, which may be due to heterogeneity within the TME. To explore this hypothesis, the spatial proximity and relationship between CD8^+^ T cells that produce IFNγ, and tumor NOS2/COX2 expressing cells was examined in 21 ER- breast tumors (including TNBC (n = 14) and HER2/neu^+^ (n = 7) phenotypes) using multiplex spatial imaging. Fluorescence imaging enables the visualization and quantification of cellular neighborhoods at the single cell level. High NOS2 and COX2 expressing cells were observed in distinct regions of the TME, as depicted in Figure 3A. Spatial distribution and density heat map analyses of the whole tumor reveal distinct regions of NOS2, COX2, and CD8^+^ T cells where NOS2 and COX2 expressing cells (Figures 3B and 3C) were observed in separate neighborhoods. In addition, higher CD8^+^ T cell densities were observed near NOS2 expressing cells while lower densities were identified near COX2-expressing clusters (Figures 3B and 3C). Thus, spatially distinct NOS2 and COX2 expressing cells in relation to CD8^+^ T cells in aggressive breast tumors suggest an association between CD8^+^ T cells and cytokine induced NOS2/COX2 niches that influence clinical outcomes.

**Figure 3.**
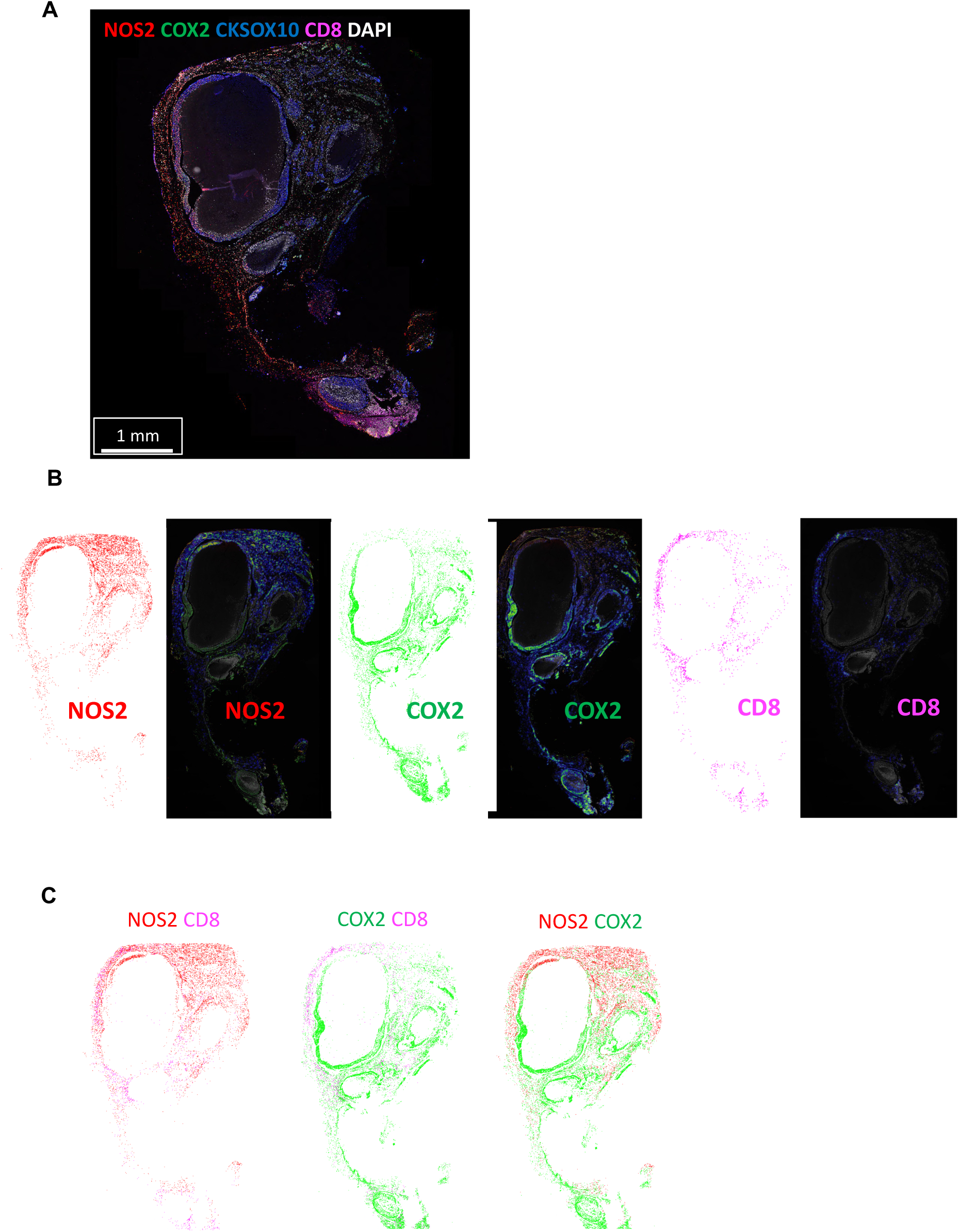
Tumor NOS2/COX2 and CD8^+^ T cells occupy unique areas in the tumor microenvironment. A) Multiplex fluorescence of an ER-/HER2+ breast tumor showing NOS2 (red), COX2 (green), CKSOX10 (blue), CD8^+^ T cells (magenta), and DAPI (white). B) spatial distribution (left) and density heat maps (right) of NOS2, COX2 or CD8+ T cells. Spatial distributions reflect a positive detection of the markers within 25µm diameter areas independent of amount. Density heat maps provide a visual quantitation reflected by color gradation (low-high) blue, green, yellow orange, and red of the biomarker protein expressions. C) comparison of NOS2/CD8, COX2/CD8, and NOS2/COX2 spatial distribution combinations.

### NOS2 and COX2 Fluorescence Intensity

The NOS2/COX2 expressions in ER- tumors previously scored as NOS2/COX2 high (hi) or low (lo) by routine immunohistochemistry (IHC) grades of 1-4 (7, 9) were analyzed for NOS2/COX2 fluorescence intensity at the single cell level using multiplexed fluorescence imaging, which provides spatial information at the single cell level in regions including necrosis, stroma, and viable tumor. Viable tumor and stroma regions were annotated by a Veterinary Pathologist on H&E images (QuPath) (22) and fused with NOS2/COX2 fluorescent expression using HALO software (Figure 4A). NOS2 and COX2 fluorescent intensities were determined for each tumor using real-time tuning in HALO software, and then mean intensities and standard deviations (SD) were determined. Threshold intensities for weak, moderate, and strong expression levels were determined by adding 2, 4, or 6 SD, respectively to the mean intensity threshold setting. NOS2/COX2 fluorescent intensities of the entire tumor quantified from these thresholds (Supplemental Figure 1A) were consistent with the original IHC Pathologist scored NOS2/COX2 expression levels previously reported (7, 9). When stratifying for tumor vs stroma, NOS2/COX2 tumor expression with strong/moderate signal intensity was significantly elevated in NOS2/COX2 high expressing tumors (Supplemental Figure 1B). In contrast, NOS2 weak signal intensities were significantly elevated in stroma but not tumor, while COX2 weak signal intensity was higher in tumor but not stroma (Supplemental Figure 1B). NOS2 and COX2 feedforward signaling (14) has been shown to maintain their expressions. A potential linear relationship between tumor NOS2 and COX2 in these tumors was examined using Pearson’s correlation coefficient, which revealed linear correlations between NOS2 and COX2 expression at strong, moderate, and weak intensities (Supplemental Figure 1C). Together, these results support NOS2/COX2 feed forward signaling as shown in Figure 1A, and as previously reported (14).

**Figure 4.**
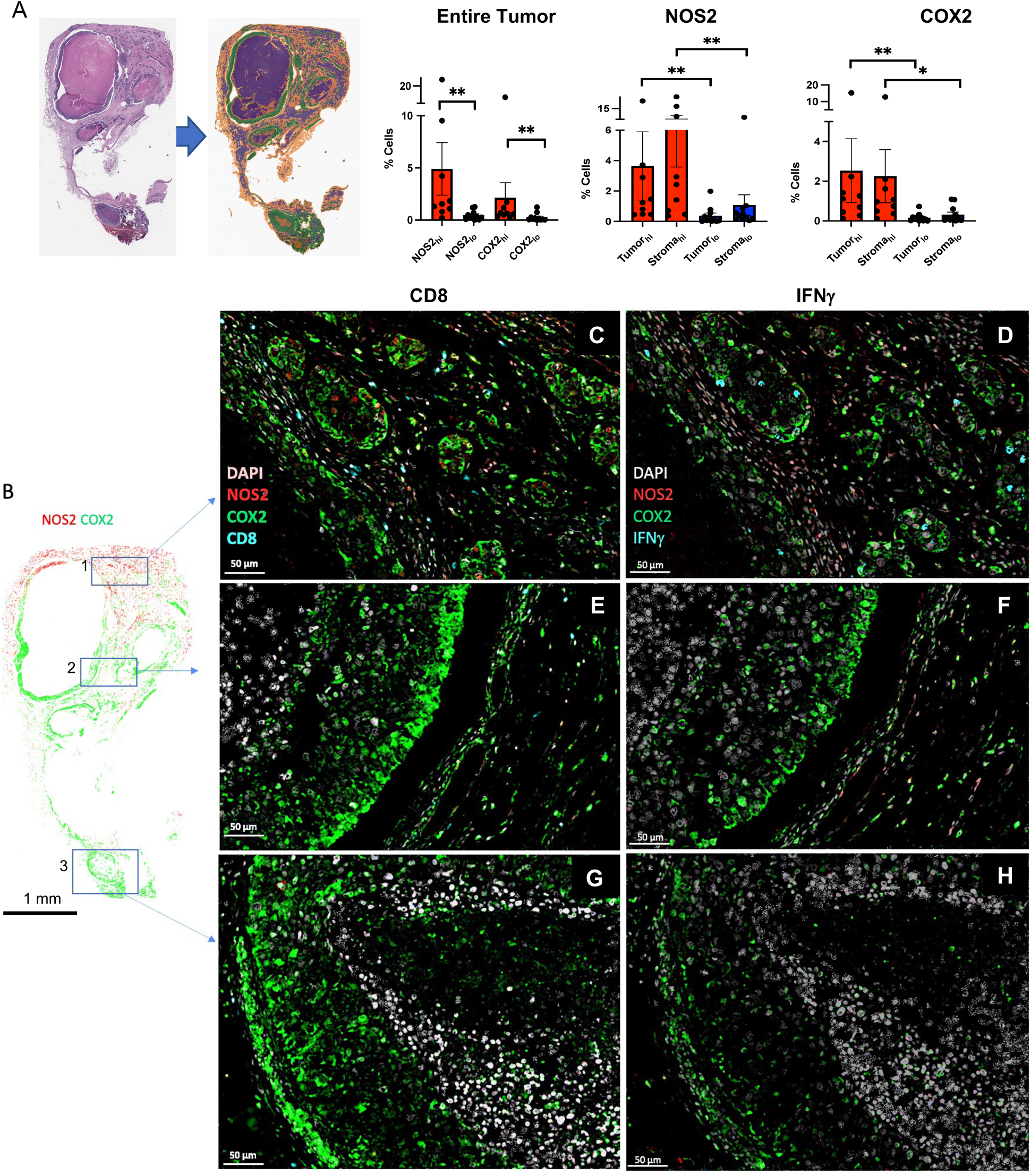
Correlation of pathology scoring and single cell fluorescence intensities. A) H&E-stained section (left) fused with a serial fluorescent image (right). H&E sections were evaluated by a pathologist who defined areas of necrosis (purple), viable tumor (green), and stroma (orange). The %NOS2/COX2 expressing cells in the entire tumor as well as tumor and stroma are shown. B) Areas in the spatial distribution highlighted by blue boxes labeled 1 (top), 2 (middle), and 3 (bottom) reflect progression from (1) inflamed regions of stroma restricted CD8^+^ T cells to (2) cold regions with reduced stroma restricted CD8^+^ T cells and (3) cold immune desert tumor core regions lacking CD8^+^ T cells. Spatial analyses of NOS2/COX2 expression in these boxed areas as well as CD8^+^ T cells or IFNγ are shown at 50μm magnification. Registered images in the inflamed region designated in box 1 show C) stroma restricted CD8^+^ T cells (cyan) or D) IFNγ expression (cyan) near NOS2 expressing cells (red). Analyses of cold regions designated in box 2 show high COX2 expression and low NOS2 expression with E) limited CD8^+^ T cells and F) limited IFNγ. Analyses of immune desert tumor core regions associated with box 3 show high tumor COX2 expression with abated levels of G) CD8+ T cells and H) abated IFNγ expression.

### Spatial Correlations of NOS2, COX2, and CD8^+^ T Cells

Figure 4A demonstrates significant increases in the %cells with elevated tumor NOS2/COX2 expression in the entire tumor as well as tumor and stroma regions. Further examination of NOS2 and COX2 spatial distributions revealed that NOS2^+^ cells are clustered at tumor margins or in stroma (Figure 4B-D). While COX2 expression was observed near NOS2 expressing cells in some regions, COX2^+^ cells were densely clustered in distinct areas deeper into the tumor core as well as in immune desert regions of the tumor (Figure 4E-H). Tumors scored as NOS2_lo_/COX2_lo_ exhibited sporadic low density NOS2 and COX2 foci, where few or no higher-intensity foci were observed. In contrast, NOS2_hi_/COX2_hi_ tumors exhibited numerous, spatially distinct, high expressing NOS2 and COX2 foci, with NOS2 clusters at the tumor-stroma interface (Figure 4C and 4D). In contrast, boxes 2-3 show COX2 clusters extending deeper into the tumor core (Figure 4E-H). Thus, NOS2 and COX2 expressing cells are spatially localized in distinct inflammatory regions of the tumor.

As depicted in Figures 1D and 2C, IFNγ is necessary for optimal NOS2/COX2 expression in MB231 cytokine treated cells. Lymphoid cells including CD8^+^ T cells are a source of IFNγsecretion (21). Recent studies have demonstrated a key role of the spatial orientation of CD8^+^ T cells for improved survival in TNBC (23). Penetration of CD8^+^ T cells into the tumor core defined a fully inflamed tumor that was predictive of improved TNBC patient survival (23). In contrast, limited CD8^+^ T cell penetration into the tumor core (≤100 CD8^+^ T cells/mm^2^) or stroma restricted CD8^+^ T cells were associated with fibrotic or immunosuppressive tumor immune microenvironments, which predicted poor survival (23). Thus, CD8^+^ T cell spatial localization is a predictor of clinical outcomes (23). Supplemental Figure 2 describes the classification of NOS2/COX2 strong single cell intensity relative to the presence of CD8^+^ T cells in all tumors. Given the predictive power of CD8^+^ T cell spatial localization (23), we observed abundant stroma restricted CD8^+^ T cells (Figure 4C) with increased IFNγ expression (Figure 4D) in regions proximal to elevated tumor NOS2 expression, indicating a potential association between CD8^+^ T cells, IFNγ, and NOS2 regulation (Figure 4C and 4D). In contrast, Figures 4E-H show areas with limited CD8^+^ T cells, as well as limited IFNγ and NOS2 expression. Importantly, COX2 is highly expressed in these regions (Figures 4E-H). Pearson’s correlation coefficients were also determined to show significant correlation between tumor NOS2_hi_ expressing cells and CD8^+^ T cells/INF*γ* (Supplementary Figure 3A). Interestingly, significant linearity was not observed between CD8^+^ T cells/IFNγ and tumor COX2 expression (Supplementary Figure 3B). These results suggest that CD8^+^ T cells could provide a source of IFNγ leading to increased tumor NOS2 expression.

### High NOS2 Cell Niches, Increased Inflammation, and Metastatic Potential

The stimulation of oncogenic pathways by NO is characterized by increased epithelial-to-mesenchymal transition (EMT), migration, and cancer cell motility culminating in cancer disease progression and metastasis (24, 25). Patients in this cohort succumbed to metastatic disease even though lymph node-positive status was *not* observed at diagnosis. NOS2_hi_ regions were near stroma restricted CD8^+^ T cells. NOS2_hi_ areas exhibited small tumor clusters that appeared to break away from the primary lesion (satellitosis), indicative of metastatic niches (Figures 5A box 1, 5B, and 5C). In contrast, satellitosis was absent in regions with lower tumor NOS2 expression as well as fewer CD8^+^ T cells and IFNγ (Figures 5D and 5E). These findings indicate that NOS2_hi_ clustering foci could promote increased metastatic potential, which is consistent with earlier reports (16, 26).

**Figure 5.**
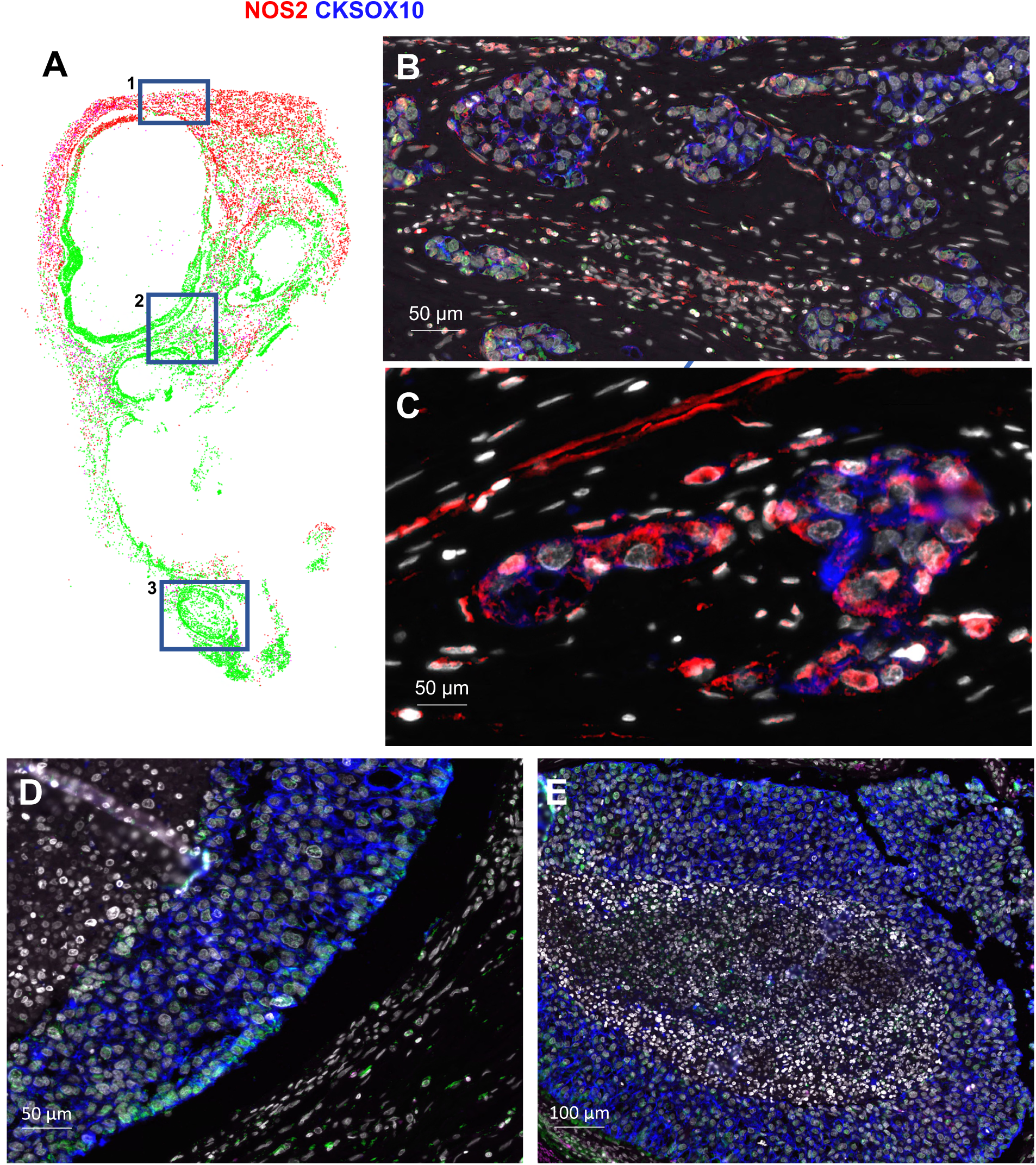
NOS2_hi_ regions are associated with increased satellitosis and metastatic potential. A) Spatial distribution of the entire tumor with blue boxes labeled 1 (top), 2 (middle), and 3 (bottom) showing B-C) the spatial localization of NOS2 (red) and the tumor marker CKSOX10 (blue) expressing cells in a NOS2_hi_ region (50μm). Both elongated and clustered NOS2 expressing cells that have broken away from the larger lesion are shown, which is indicative of satellitosis and increased metastatic potential. These phenotypes are not observed in cold immune regions (D) or immune desert regions (E).

### Morphological Changes that Mimic NOS2^+^ Niches

Cellular morphology is a key aspect of metastasis, where cells acquire an elongated phenotype during migration and invasion processes. *In vitro* migration models have shown NO roles during metastatic processes, where exposure to higher NO flux (100–300 nM) for 24-48 hr increased *in vitro* migration and invasion of MB231 and MB468 breast cancer cells (7, 16). These earlier observations suggest that increased tumor NOS2 expression and clustering would generate a flux of NO that enhances the metastatic potential of exposed cells within that niche (16). Herein, we further explored the influence of tumor cell NOS2/COX2 expression on altered cellular morphology. As shown in Figure 6A and 6B, MB231 cells exposed to individual, or combination cytokine treatment demonstrated morphological changes and cellular elongation characteristic of EMT in migrating and invading cells. In addition, scratch test assay showed increased wound closure after 12 hr of IFNγ + TNFα combination treatment when compared to the untreated control cells (Figure 6C). Similarly, Boyden chamber assays showed increased cell invasion in response to 48 hr IFNγ + TNFα combination treatment, which was reduced by the pan-NOS/COX inhibitors (LNAME/Indomethacin) both as single treatment agents and in combination (Figure 6D). These results suggest that the upregulation of NOS2/COX2 tumor expression within an inflammatory niche could generate phenotypes with increased metastatic potential (Figure 6).

**Figure 6.**
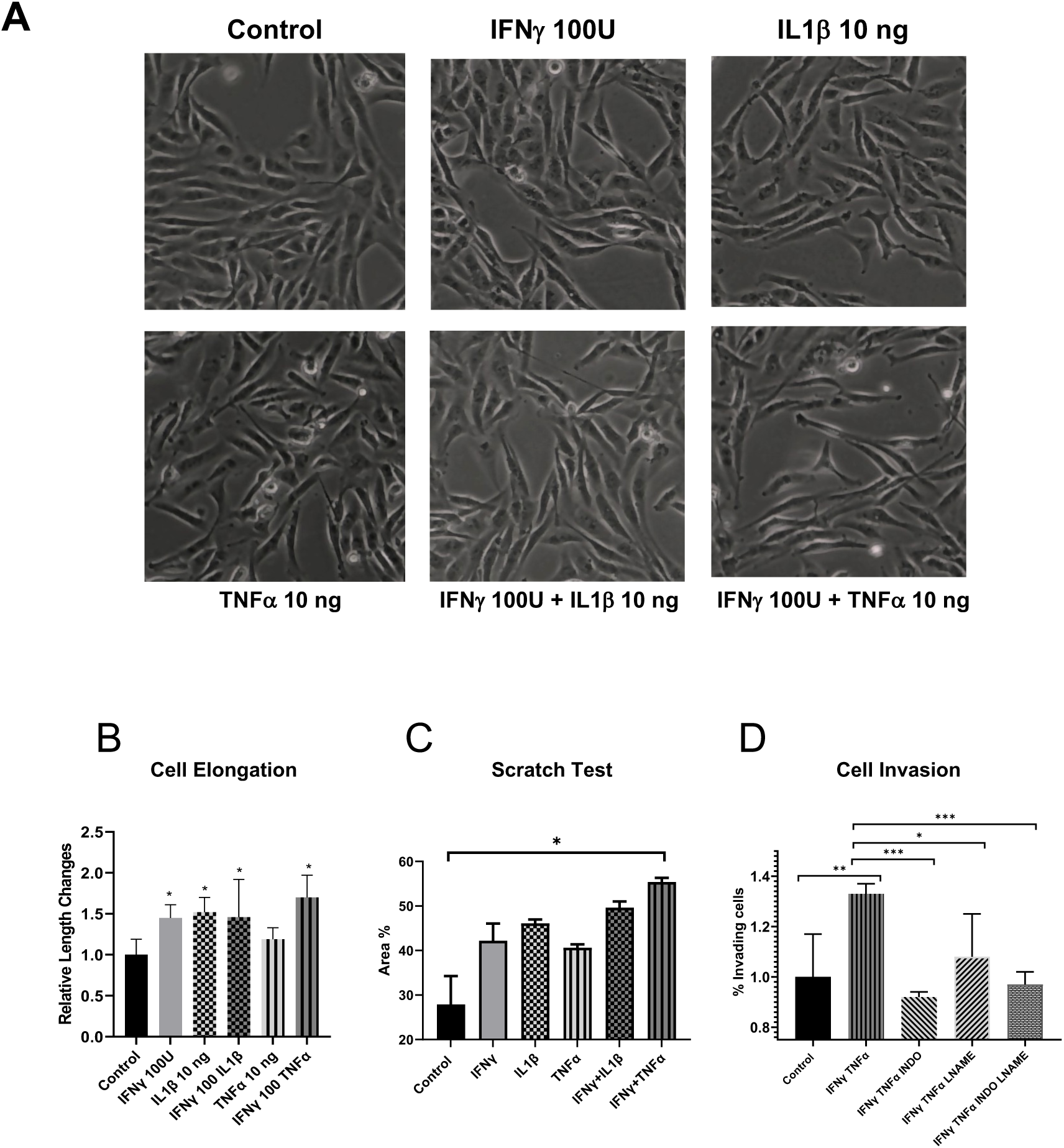
IFNγ and Cytokines promote elongation and migration MB231 breast cancer cells. A) Microscope images (10X) of control and cytokine treated MB231 cells for 48 hr. B) Cell elongation analysis after 48 hr cytokine treatment. C) A scratch test assay of MB231 cells after 12 hr of cytokine treatment. D) Cell invasion assay; MB231 cells were seeded in the upper well of a Boyden chamber with serum-free media ± cytokines and the pan-NOS/COX inhibitors LNAME and indomethacin for 48 hr. Lower chamber was filled with complete media. Cells were counted against the standard curve. Results are presented as mean ± SD. * p < 0.05, ** p < 0.001, *** p < 0.0001.

## Discussion

One of the most effective prognostic indicators for ER- breast cancer is the association between NOS2 and COX2 (14, 27). The data above demonstrates an unusual link between IFNγ and lymphoid cells in terms of clinical outcomes as summarized in Figure 7. IFNγand CD8^+^ T cells are linked to and necessary for the production of high NOS2 and COX2 levels, despite the fact that they are predictive of positive clinical outcomes and a hallmark of a good prognosis in many malignancies (23, 28). According to the scRNAseq results, IFNγ and IL1β/TNFα are necessary to induce the highest levels of NOS2 and COX2 expression, which suggests that lymphoid cells must be a contributing factor in the tumor. According to earlier research, IFNγ is critical for the stimulation of NOS2 in DLD1 (human colon cancer) cells (29). Also, the scRNAseq of MB231 with IFNγ+IL1β reveals a strong connection between NOS2/COX2, and IL1β, TNFα, IL6, and IL8, bolstering the notion of a strengthened Th1 microenvironment acquired from the TCGA (Figure 1). This implicates multifactor immune mechanisms leading to a feedforward loop that promotes the induction of high NOS2/COX2 expressing cellular niches and disease progression. Previous studies have demonstrated that IFNγ, IL6, PGE2, and IL1 all boosted NOS2 expression, suggesting that several reinforcing processes were involved in NOS2 upregulation (16). In addition, IL1α is a mediator of ER stress that is frequently observed in the TME after chemotherapy. Both IL1αand ILβ increased in response to IFNγand IL1β significantly enhances these complimentary pathways for sustained elevation of NOS2/COX2 mechanisms. These factors could conspire to create a NOS2/COX2 inflammatory niche that promotes disease progression.

**Figure 7.**
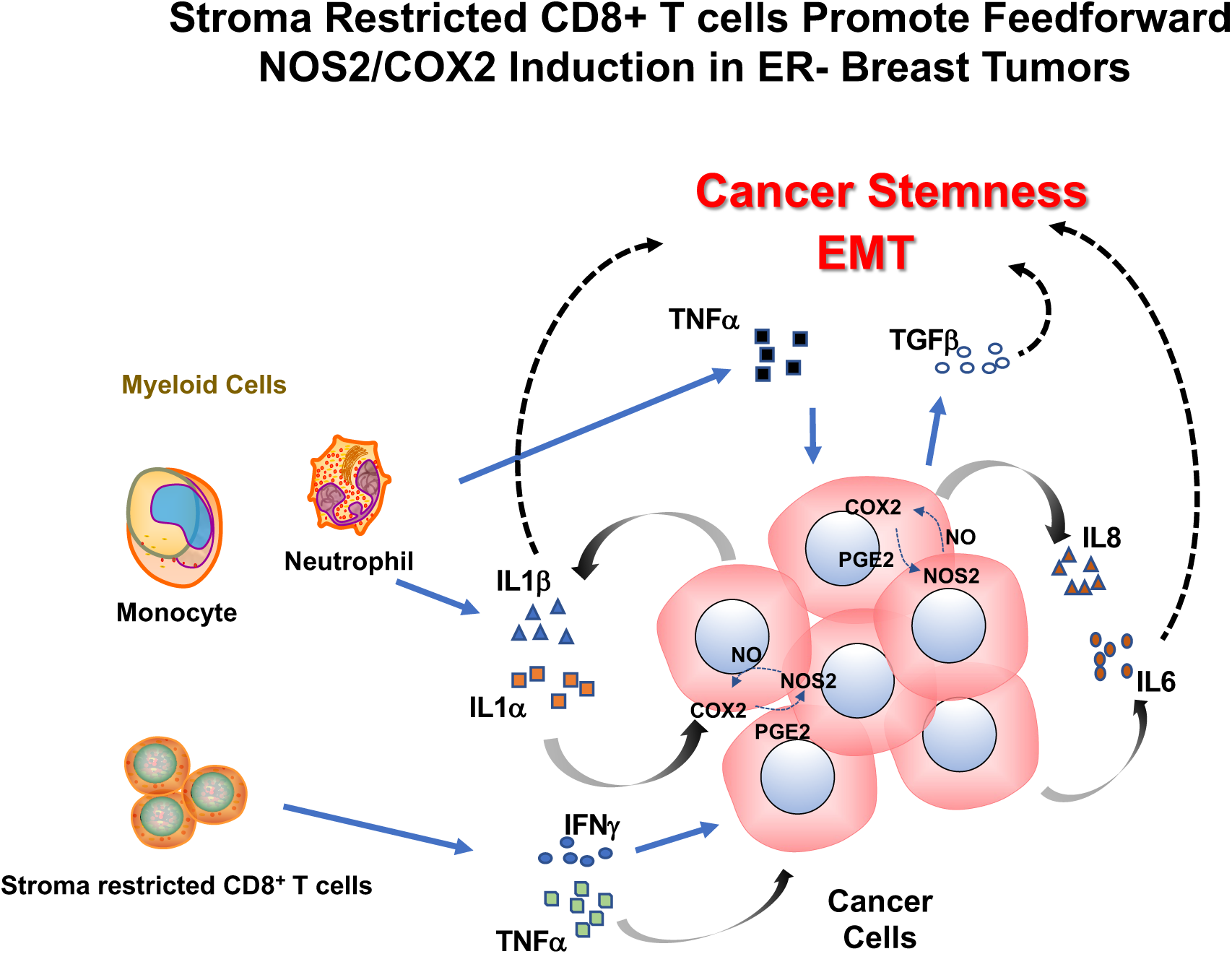
Interplay of cytokine production in the tumor microenvironment leading to NOS2_hi_/COX2_hi_ tumor expressing regions. The secretion of IFNγ by stroma restricted CD8^+^ Tcells and IL1β/TNFα secreted by myeloid cells within the tumor microenvironment leads to tumor NOS2/COX2 expression and the development of aggressive cellular niches with increased metastatic potential and promote immunosuppression. Cellular neighborhoods expressing high tumor NOS2/COX2 then increase IL1α/β creating a feedforward loop that maintains tuor NOS2/COX2 expression and elevated cytokines including IL-8 and IL-6 as well as the activation of latent TGFβby NO. These factors conspire to promote immunosuppression, metastasis, and cancer stemness through NOS2-derived NO and COX2-derived PGE2.

### Single Cell Implications of NOS2 Expression

Despite the protein’s sequence and biochemistry being comparable in the mouse and human, the NOS2 promoter is complicated and differs significantly between the two species (29–31). Expression of NOS2 in murine macrophages maximally induced *in vitro* by IFNγ or LPS has been the gold standard in the NO field, where estimated NO flux could reach as high as 1-5 µM at the cellular level (13). Murine tumor cells displayed significantly higher NOS2 activity in response to IFNγ or LPS, suggesting a key distinction between tumor cells and macrophages (13). While data here clearly demonstrate that maximum human NOS2 expression is induced by IFNγ, IL1β, and TNFα, identical to that in murine tumor cells, human macrophages do not activate NOS2 with IFNγ or LPS. However, the levels of NO produced by mouse and human tumor cells still differ significantly on a fundamental level. Recently, it was demonstrated that the number of NOS2^+^ cells, rather than NOS2 expression, correlated with the amount of NO and nitrite produced *in vitro* (11). As a result, the clusters of NOS2 expressing cells will affect NO levels, and NOS2 cell clustering can produce areas of greater NO flux (13, 32). *In vitro* experiments reveal that high nitrite, and NO levels are present when 50–80% of the cells express NOS2 and fluxes are higher than 100 nM. However, NO production is an order of magnitude lower in 5% of human cancers. Our earlier research demonstrates that NO-driven carcinogenic pathways take place at an ideal concentration of 200-400 nM, which increases the expression of IL6 and IL8 (12). Nonetheless, NO levels are higher where NOS2-expressing cells are concentrated at a much higher density, such as in localized foci inside the tumor. When considered collectively, these findings suggest that these regions of high-density NOS2-expressing cells have larger NO flux, which can trigger oncogenic mechanisms that take place in the range of 100-300 nM NO in the petri dish (33–35).

### The Spatial Configuration of the NOS2/COX2 Niche

Areas enriched for CD8^+^ T cells and IFNγ are related to the juxtaposition of lymphoid and tumor cells within the TME, which results in high clustering of enhanced NOS2 expressing cells. An earlier study looking at CD8^+^ T cell placement demonstrated that spatial orientation is a crucial factor in the determination of TNBC clinical outcomes (23). Positive results are described by tumor penetrating CD8^+^ T cells deep into the tumor core in a completely inflamed tumor. On the other hand, stroma restricted CD8^+^ T cells and immune desert regions devoid of CD8^+^ T cells predict poor clinical outcomes (23). Here we demonstrate that high NOS2 cellular niches can be formed at the tumor margin and proximal to stroma restricted CD8^+^ T cells. These niches can now encounter IFNγ and other cytokines that induce tumor NOS2/COX2 expression. Contrarily, NOS2^+^ and COX2^+^ cells are scattered and observed at lower levels in areas with increased CD8^+^ T cell penetration into the tumor. The aforementioned information demonstrates unequivocally that CD8^+^ T cell and IFNγ are close to NOS2 and suggests that an inflammatory niche with stroma restricted CD8^+^ T cells is necessary for NOS2 induction.

An immune desert devoid of CD8^+^ T cells is another significant aspect of the TME that has previously been identified (23). Poor clinical outcome is suggested by low CD8^+^ T cell counts and exhausted CD8^+^ T cells in the tumor compartment (36). One of the critical factors in the absence of CD8^+^ T cells associated with low IFNγin immunological desert regions was also revealed in regions deficient in NOS2 with increased COX2 expression. This implies that the immunological desert is associated with COX2 positive and NOS2 negative regions. Elevated CD8^+^ T cells and other lymphoid cells that are restricted to tumor stroma or margins can result in situations that promote higher NOS2 and COX2.

### Ramification of NOS2/COX2 Niche and Metastatic Potential

Our prior research demonstrated that NO is essential for promoting EMT and metastasis (16). Increased inflammation at these NOS2 foci increases the likelihood for metastasis, and the discovery of the NOS2 positive niche at the tumor stroma interface suggests that this may be the site of metastasis. Elongation and EMT induced by NO are known to mediate these effects (16). Also, IL1 and PGE2 enhance EMT and cell motility in breast cancer (37, 38). Herein, IFNγ and ILβ1/TNFα promote motility and elongation (7). As a result, the NOS2/COX2 inflammatory niches increase the potential for cancer cell motility and metastatic spread. Therefore, limited metastatic potential can be achieved by inhibition of NOS2/COX2 feedforward loops (39). Given that metastasis is the primary cause of cancer deaths, NOS2/COX2 spatial localization at these sites of inflammation could provide an early prognostic indicator of poor outcome even in the absence of lymph node positive status (7).

## Summary

IFNγ plays a key role in the induction of proinflammatory antitumor immune responses (40). However, recent studies have shown that IFNγ response is concentration dependent where low levels in the TME promote protumorigenic disease progression mediated in part through the downregulation of major histocompatibility complexes and upregulation of indoleamine 2,3-dioxygenase and programmed cell death ligand 1 (40). In addition, IFNγ is necessary to stimulate tumor specific NOS2/COX2 expression, which through a multifaceted process also drives oncogenic pathways and shapes immunological profiles associated with poor prognosis (10, 11). Given that IFNγ is secreted by cytolytic CD8^+^ T cells, spatial analysis suggests that the quantity and location of CD8^+^ T cells (23) present an opportunity for the formation of IFNγ regulatory processes within the TME, including the upregulation of tumor NOS2/COX2 expression and the development of niches that promote disease progression, metastasis, and poor clinical outcomes (7, 10, 11, 23).

## Materials and Methods

## Cell Culture

The MDA-MB231 (MB231) human breast cancer cell line was obtained from the American Type Culture Collection (ATCC, Manassas, VA) and grown in RPM1-1640 (Invitrogen) supplemented with 10% fetal bovine serum (FBS; Invitrogen, Waltham, MA) at 37°C in a humidified atmosphere of 5% CO_2_ in air. Cells were serum starved overnight prior to experimentation. Depending on the downstream assays, the cells were incubated for 12, 24, or 48 hr with the addition of ddH2O (control), IFNγ 100 U/mL (285-IF/CF, R&D Systems, Minneapolis, MN), IL1β 10 ng/mL (201-LB/CF, R&D Systems), TNFα 10 ng/mL (210-TA/CF, R&D Systems), IL17A 10 ng/mL (7955-IL-CF, R&D Systems), lipopolysaccharide (LPS, Sigma, St. Louis, MO) 10 mg/mL (L2630, Sigma), LNAME 500 mM (N5751, Sigma), and/or Indomethacin 100 μM (I7378, Sigma).

### *In Vitro* Scratch Assay

One million cells were plated in a 60-mm dish and allowed to reach 100% confluency. A 200 μl pipette tip was used to etch a straight scratch line across the confluent monolayer. Floating and dead cells were eliminated by washing the dishes in 1X PBS and then complete media was added. A 10x objective inverted microscope (EVOS, Life Technologies, Carlsbad, CA) was used to take images at 0, 4, 8, and 12 hr. The opensource software ImageJ measures the pace at which scratch gaps refilled (version 1.53u) (41).

### Cell Invasion Assay

A cell invasion assay (cat# ab235697) in 96-well plate format from Abcam (Waltham, MA) was used. After cell synchronization, complete medium was given to the lower chamber as an attractant, and 50,000 cells were seeded in the upper chamber with cytokines +/- inhibitors for 48 hr. Migrated fluorescent cells were counted at Ex/EM = 530/590 nm on a SpectraMax i3x plate reader (Molecular Devices, San Jose, CA) and compared to a standard curve made from the same cell line.

### Single Cell RNAseq

The single cell library was generated using the 10x Genomics (San Francisco, CA) Single Cell 3’ Reagent Kit v3 and then sequenced in our sequencing facility (NCI at Frederick, MD) using an Illumina NovaSeq 6000. Sample cells in suspension medium were examined for viability before library preparation. The cDNAs were sequenced after being barcoded, pooled, and amplified during the library preparation. On average, 10,000 cells per sample were sequenced. The Cell Ranger software provided raw reads as input (10x Genomics, Version 6.1.2). They were demultiplexed and converted into BCL files using the Cell Ranger. All readings were mapped to the human reference genome using the default 10x Genomics Pipeline (Version 3.1.0) after passing quality checks (GRCH38-30.0). Annotated transcript counts within each cell were used to construct UMI (Unique Molecular Identifier) count matrices.

### scRNAseq Data Analysis

The matrix h5 files for each sample were uploaded to the internal Partek (St. Louis, MO) Flow server for data processing and data mining. All counts were normalized using the default "counts per million, add 1, and log2 transformed" method. The GSA (Gene Specific Analysis) tool was then applied to discover genes that were differentially expressed between the various experimental samples. Absolute fold changes ≥ 2 and a p value < 0.05 were used to select genes. We utilized the Loupe Browser (version 6.3.0, 10x Genomics) to visually inspect the aggregated and standalone datasets to analyze the clustering patterns of the scRNAseq data. To create correlation heat maps, scRNAseq datasets that had been directly processed in Seurat (version 4.0, Satija lab, https://satijalab.org/seurat/) from the Cell Ranger output were exported to RStudio (2022.07.2 Build 576, https://posit.co/) in parallel. Single cell data is available upon request to the Corresponding Author.

### The Cancer Genome Atlas (TCGA) Analysis

The breast cancer (BRCA) subset of TCGA (https://www.cancer.gov/about-nci/organization/ccg/research/structural-genomics/tcga) were accessed through the UCSC (University of California, Santa Cruz) Xena Browser (Date of access: 11/2/2022, https://xena.ucsc.edu/). In brief, all targeted Th1, Th2, Th17 cytokines, NOS2 and COX2 genes were surveyed in Xena browser, and then exported to RStudio (2022.07.2 Build 576) for subsequent correlation analysis.

### Tissue Collection and Immunohistochemical Analysis of Patient Tumor Sections

Tumor specimens (n = 21) were obtained from breast cancer patients recruited at the University of Maryland (UMD) Medical Center, the Baltimore Veterans Affairs Medical Center, Union Memorial Hospital, Mercy Medical Center, and the Sinai Hospital in Baltimore between 1993 and 2003. Informed consent was obtained from all patients. The collection of tumor specimens, survey data, and clinical and pathological information (UMD protocol no. 0298229) was reviewed and approved by the UMD Institutional Review Board (IRB) for the participating institutions. The research was also reviewed and approved by the NIH Office of Human Subjects Research (OHSR no. 2248). Breast tumor NOS2 and COX2 expression was analyzed previously by IHC using 1:250 diluted NOS2 antibody and 1:50 diluted COX2 antibody (no. 610328 and 610204, respectively, BD Biosciences, San Diego, CA,) and scored by a pathologist (7, 9). For NOS2 staining, a combination score of intensity and distribution were used to categorize the immunohistochemical NOS2 stains where intensity received a score of 0-3 if the staining was negative, weak, moderate, or strong. The NOS2 distribution received scores of 0-4 for distributions <10%, 10-30%, >30-50%, >50-80% and >80% positive cells (7). For COX2 staining, scores of negative to weak (1–2) or moderate to strong (3–4) were categorized as low or high, respectively (9). Herein, NOS2 and COX2 expressions were also analyzed by fluorescent staining performed on the Leica Biosystems (Wetzlar, Germany) Bond RX Autostainer XL ST5010 using the Bond Polymer Refine Kit (Leica Biosystems DS9800), with omission of the Post Primary Block reagent, DAB and Hematoxylin. After antigen retrieval with EDTA (Bond Epitope Retrieval 2), sections were incubated for 30 min with COX2 (Cell Signaling Technology, Danvers, MA, no. 12282, 1:100), followed by the Polymer reagent and OPAL Fluorophore 520 (AKOYA, Marlborough, MA). The COX2 antibody complex was stripped by heating with Bond Epitope Retrieval 2. Sections were then incubated for 30 min with NOS2 antibody (Abcam no. ab15323, 1:50), followed by the Polymer reagent and OPAL Fluorophore 690. The NOS2 antibody complex was stripped by heating with Bond Epitope Retrieval 2 and then stained with CD8 (Abcam no. 101500, 1:100) or IFNγ (Abcam no. 231036, 1:200), followed by the Polymer reagent and OPAL Fluorophore 570. Sections were stained with DAPI and coverslipped with Prolong Gold AntiFade Reagent (Invitrogen). Images were captured using the Aperio ScanScope FL whole slide scanner (Leica). The original IHC previously reported (7, 9) and fluorescent NOS2/COX2 staining results were generally consistent.

Formalin-fixed paraffin embedded (FFPE) tissue sectioned at 4 μm and mounted on SuperFrost Plus slides were stained with a FixVUE Immuno-8^TM^ Kit (formerly referred to as UltiMapper® kits (Ultivue Inc., Cambridge, MA), USA; CD8, NOS2, COX2, CKSOX10, and IFNγ cocktail) using the antibody conjugated DNA-barcoded multiplexed immunofluorescence (mIF) method (1). These kits include the required buffers and reagents to run the assays: antibody diluent, pre-amplification mix, amplification enzyme and buffer, fluorescent probes and corresponding buffer, and nuclear counterstain reagent. Hematoxylin and Eosin (H&E) and mIF staining was performed using the Leica Biosystems BOND RX Autostainer. Before performing the mIF staining, FFPE tissue sections were baked vertically at 60-65 °C for 30 min to remove excess paraffin prior to loading on the BOND RX. The BOND RX was used to stain the slides with the recommended FixVUE (UltiMapper) protocol. During assay setup, the reagents from the kit were prepared and loaded onto the Autostainer in Leica Titration containers. Solutions for epitope retrieval (ER2, Leica Biosystems cat# AR9640), BOND Wash (Leica Biosystems cat# AR9590), along with all other BOND RX bulk reagents were purchased from Leica). During this assay, the sample was first incubated with a mixture of all 4 antibody conjugates, next the DNA barcodes of each target were simultaneously amplified to improve the sensitivity of the assay. Fluorescent probes conjugated with complementary DNA barcodes were then added to the sample to bind and label the targets; Next, a gentle signal removal step was used to remove the fluorescent probes of the markers. The stained slides were mounted in Prolong Gold Anti-Fade mountant (Thermo Fisher Scientific, Waltham, MA, cat# P36965 and coverslipped (Fisherbrand Cover Glass 22 x 40mm, #1.5). Digital immunofluorescence images were scanned at 20× magnification. Images were co-registered and stacked with Ultivue UltiStacker software. The digital images were then analyzed using HALO image analysis platform (42).

### Statistical Analysis

Experiments were assayed in triplicate unless otherwise stated. Student t test was test was employed to assess statistical significance using the GraphPad Prism software (version 9). Image analyses are reported as mean ± SEM and T tests with Welch’s or Mann Whitney correction were used when appropriate to determine significance. Linear analyses and Pearson’s correlations were also conducted to determine significant correlations between protein expressions using Prism software. Significance is reported as *p ≤ 0.05, **p ≤ 0.01, ***p ≤ 0.001, ****p≤0.0001. Single cell correlation analyses were conducted in RStudio using the corrplot (0.92) in R (4.2.1).

## Acknowledgements

This project was funded in whole or in part with Federal funds from the Intramural Research Program of the NIH, National Cancer Institute, CCR, CIL (RYSC, LAR, AG, EF, HR, VS, RJK, SMH, DWM, SKA, SA, DAW). This research was supported in part by the Intramural Research Program of NIH, Frederick National Lab, Center for Cancer Research 75N91019D00024 (ALW, WFH, MP, SKA, SJL), NIH R01CA238727, NIH U01CA253553, and John S Dunn Research Foundation (STCW), NCI grant no. U54 CA210181, the Breast Cancer Research Foundation (BCRF), the Moran Foundation, Causes for a Cure, philanthropic support from M. Neal and R. Neal, and the Center for Drug Repositioning and Development Program (CREDO) (JCC), Science Foundation Ireland (SFI) grant number 17/CDA/4638, and a SFI and European Regional Development Fund (ERDF) grant number 13/RC/2073 (SAG). The content of this publication does not necessarily reflect the views or policies of the Department of Health and Human Services, nor does mention of trade names, commercial products, or organizations imply endorsement by the US Government.

## Abbreviations

BRCA: Breast cancer
CKSOX10: pan-cytokeratin/ Transcription factor
SOX10: COX2 Cyclooxygenase 2 (official name: PTGS2)
COX2hi: COX2 high
COX2lo: COX2 low
DAPI: 4′,6-diamidino-2-phenylindole
EMT: Epithelial-to-mesenchymal transition
ER: Endoplasmic reticulum
ER-: Estrogen receptor negative
ER+: Estrogen receptor positive
GSA: Gene Specific Analysis
HER2/HER2/neu: Human epidermal growth factor 2
IFNγ: Interferon gamma
IHC: Immunohistochemistry
IL1: Interleukin 1
IL1α: Interleukin 1 alpha
IL1β: Interleukin 1 beta
IL17/IL17A: Interleukin 17/Interleukin 17A
IL6: Interleukin 6
IL8: Interleukin 8
IL10: Interleukin 10
Indo: Indomethacin
IRB: Institutional Review Board
LNAME: Nitro-L-arginine methyl ester hydrochloride
LPS: Lipopolysaccharide
MB231: MDA-MB231
NO: Nitric oxide
NOS2: Nitric oxide synthase 2
NOS2+: NOS2 positive cell
NOS2-: NOS2 negative cell
NOS2hi: NOS2 high
NOS2lo: NOS2 low
PGE2: Prostaglandin E2
PTGS2: Prostaglandin-endoperoxide synthase 2
scRNAseq: Single-cell RNA sequencing
t-SNE: t-distributed stochastic neighbor embedding
TCGA: The cancer genome atlas
Th1: T helper type 1
Th2: T helper type 2
Th17: T helper type 17
TIMP1: TIMP metallopeptidase inhibitor 1
TLR4: Tool-like receptor 4
TME: Tumor microenvironment
TNFα/TNF: Tumor Necrosis Factor Alpha
UMD: University of Maryland
UMI: Unique Molecular Identifier
TNBC: Triple negative breast cancer

---

**Supplemental Figure 1.**
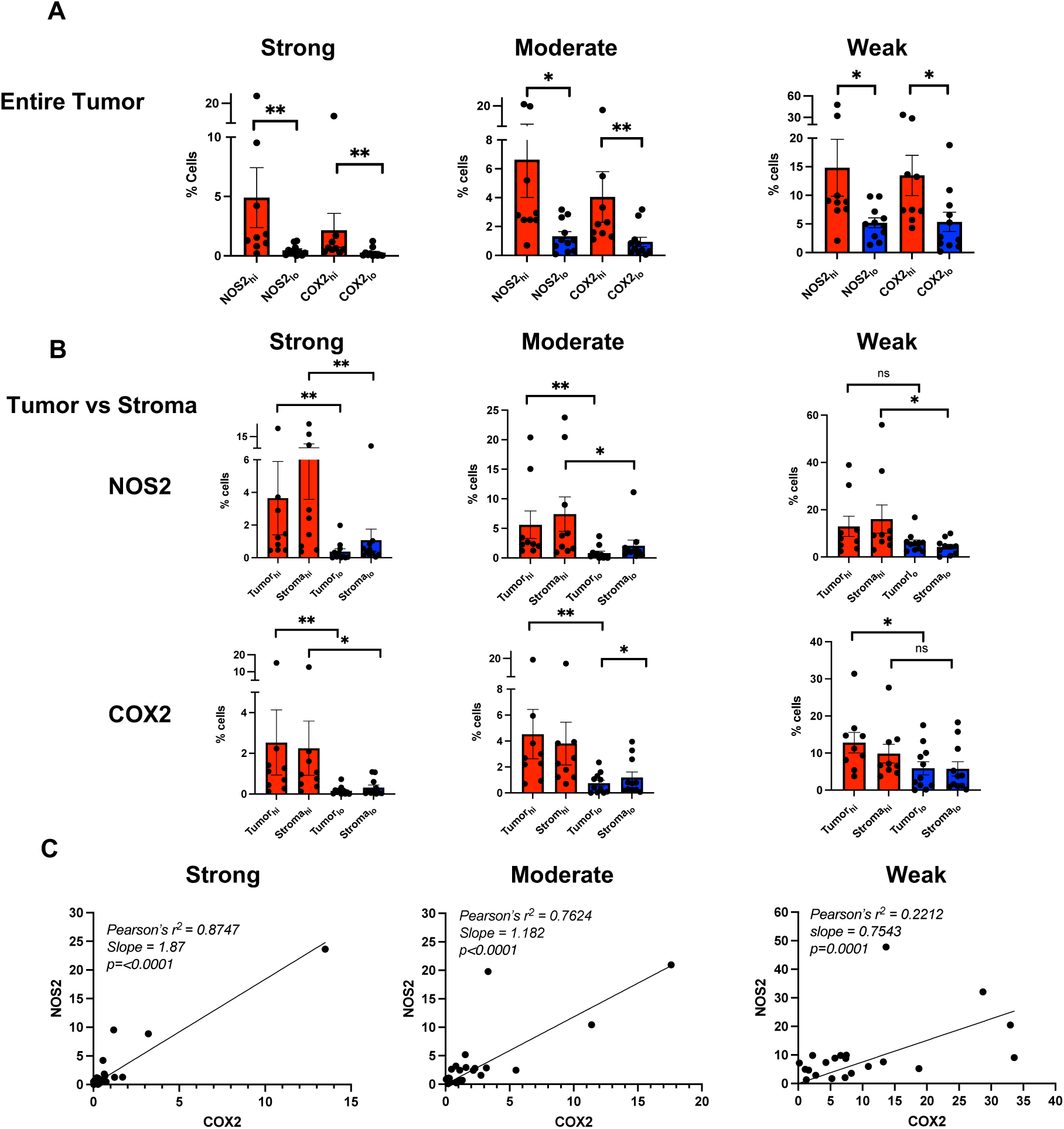
N**O**S2 **and COX2 spatial localization and signal intensity.** A) The entire tumor was analyzed for strong, moderate, and weak NOS2 and COX2 fluorescence intensities at the single cell level in 21 NOS2_hi_/COX2_hi_ (n=10) and NOS2_lo_/ COX2_lo_ (n=11) tumor images and found to be consistent with the original IHC Pathologist scoring for area and intensity as reported (7, 9). B) Analysis of strong, moderate, and weak NOS2/COX2 signal intensities in Tumor vs Stroma regions are presented. NOS2 and COX2 fluorescent intensities were determined for each tumor using real-time tuning in HALO software, and then mean intensities and standard deviations (SD) were determined. Threshold intensities for weak, moderate, and strong expression levels were determined by adding 2, 4, or 6 SD, respectively to the mean intensity threshold setting. C) Pearson’s correlation coefficients and linear regression analyses were performed to determine statistically significant linear relationships for NOS2 vs COX2 tumor expression at strong, moderate, and weak signal intensities.

**Supplemental Figure 2.**
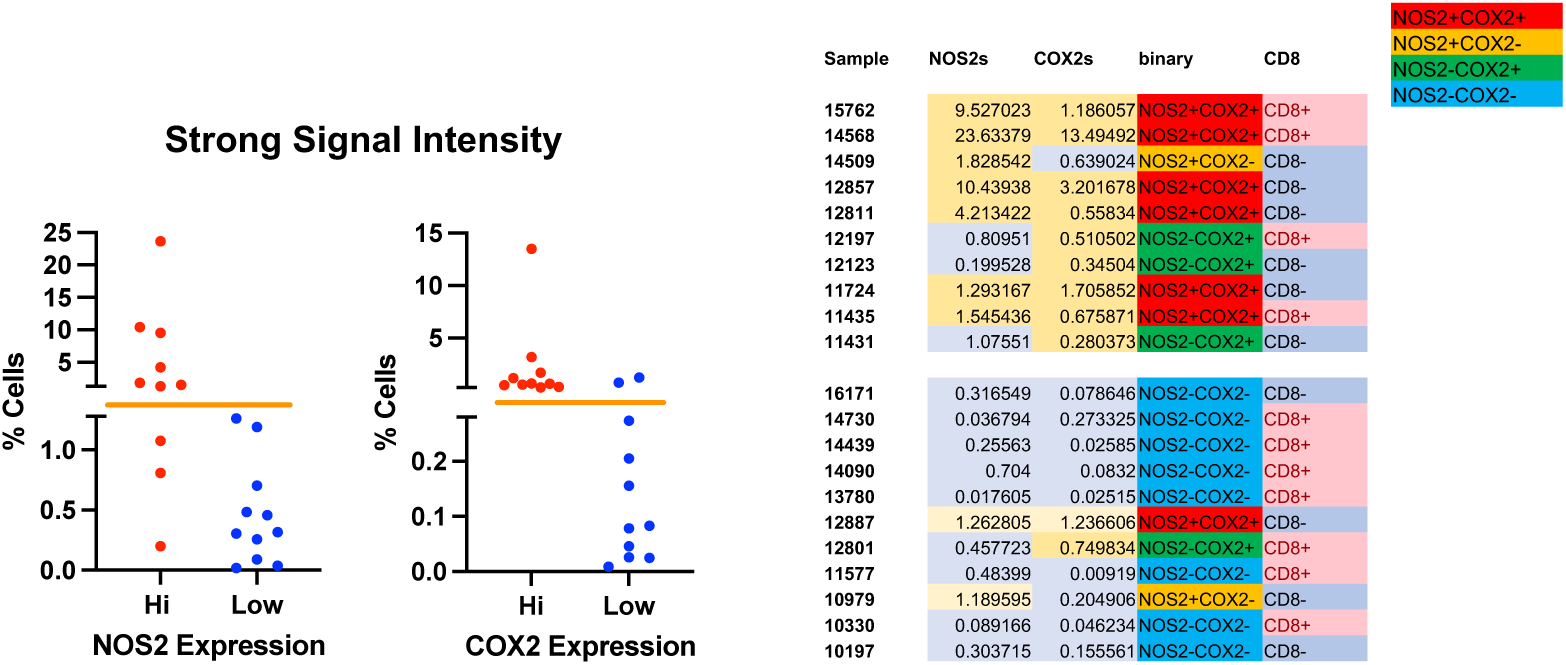
Classification of each tumor based on the single cell level of strong intensity NOS2/COX2 expression. The original IHC’s were designated NOS2/COX2 hi or low based upon Pathologist scoring of area and intensity of the entire tumor. Herein, the selection of population distribution between low and hi was used to determine NOS2/COX2 classification. These hi and low NOS2/COX2 population distributions were assigned as +/- to classify tumors based on the single cell NOS2/COX2 strongest thresholds and expressing cells as percentage of the total. The designation of CD8^+^ T cells was determined as above or below the mean of 5%.

**Supplemental Figure 3.**
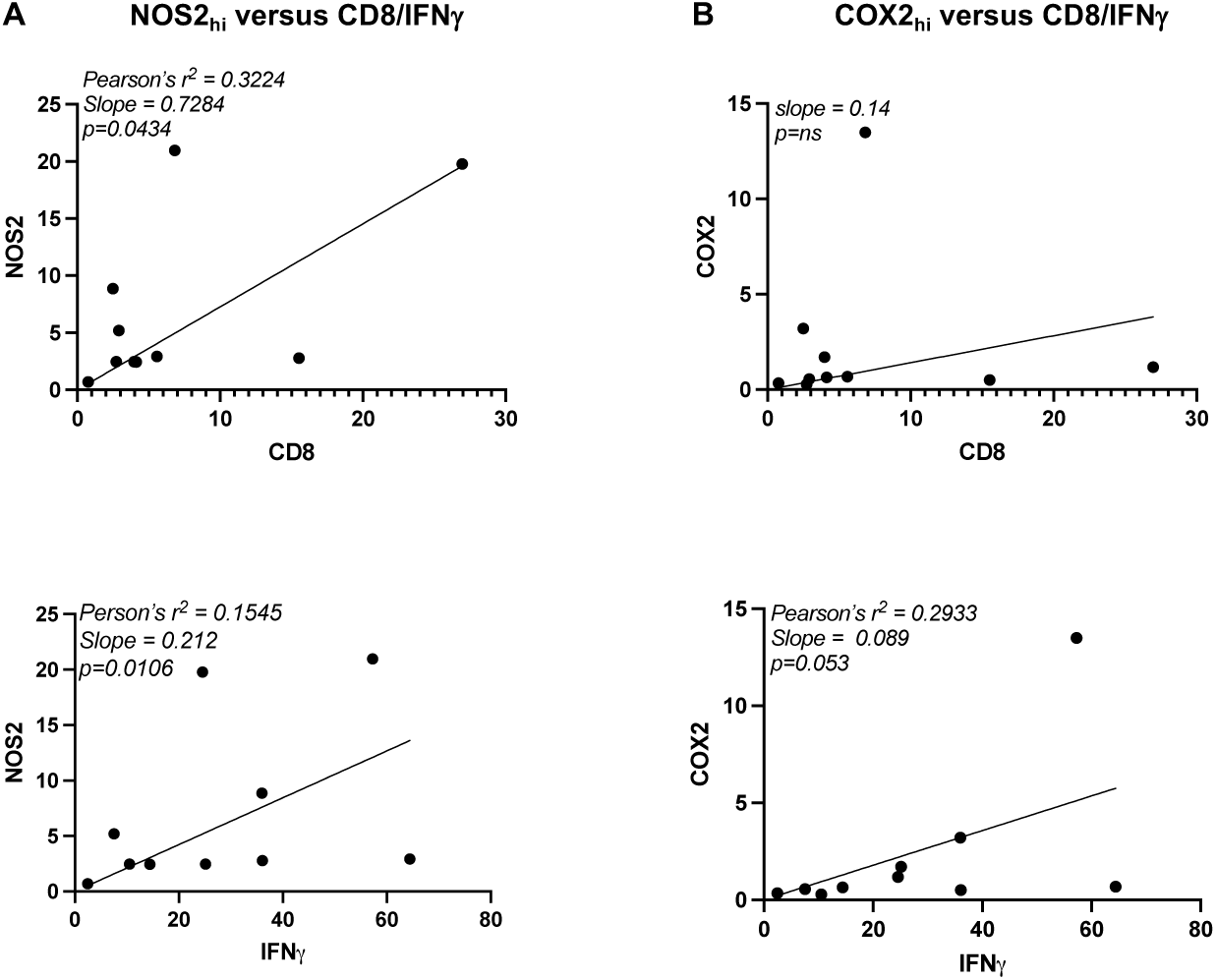
L**i**near **correlation analyses.** Pearson’s correlation coefficients and linear regression analyses were performed to determine statistically significant linear relationships between Tumor NOS2^+^ vs CD8^+^ T cells/IFNγ (A) and Tumor COX2^+^ vs CD8^+^ T cells/IFNγ (B) in areas stratified for high intensity tumor NOS2/COX2 expression. Pearson’s r^2^ correlations, slopes and p values were determined and shown in each graph.

